# Hypothetical Human Immune Genome Complex Gradient May Help to Explain the Congenital Zika Symdrome Catastrophe in Brazil: A New Theory

**DOI:** 10.1101/2020.06.03.132878

**Authors:** F.R.S Oliveira, C.C. Bresani-Salvi, C. N.L. Morais, A.W. Bigham, U. Braga-Neto, G.E. Maestre, j. l. Wandebergh, E.T.A Marques, B. Acioli-Santos

## Abstract

There are few data considering human genetics as an important risk factor for birth abnormalities related to ZIKV infection during pregnancy, even though sub-Saharan African populations are apparently more resistant to CZS as compared to populations in the Americas. We hypothesized that single nucleotide variants (SNVs), especially in innate immune genes, could make some populations more susceptible to Zika congenital complications than others. Differences in the SNV frequencies among continental populations provide great potential for Machine Learning techniques. We explored a key immune genomic gradient between individuals from Africa, Asia and Latin America, working with complex signatures, using 297 SNVs. We employed a two-step approach. In the first step, decision trees (DTs) were used to extract the most discriminating SNVs among populations. In the second step, machine learning algorithms were used to evaluate the quality of the SNV pool identified in step one for discriminating between individuals from sub-Saharan African and Latin-American populations. Our results suggest that 10 SNVs from 10 genes (CLEC4M, CD58, OAS2, CD80, VEPH1, CTLA4, CD274, CD209, PLAAT4, CREB3L1) were able to discriminate sub-Saharan Africans from Latin American populations using only immune genome data, with an accuracy close to 100%. Moreover, we found that these SNVs form a genome gradient across the three main continental populations. These SNVs are important elements of the innate immune system and in the response against viruses. Our data support the *Human Immune Genome Complex Gradient* hypothesis as a new theory that may help to explain the CZS catastrophe in Brazil.

## INTRODUCTION

Zika virus (ZIKV), an arbovirus that belongs to the *Flaviviridae* family, was initially isolated from a captive rhesus monkey in Uganda in 1947 (Dick et al. 1952; Dick et al. 1953). The most common reported ZIKV infection symptoms in humans are rash, fever, arthralgia, and conjunctivitis. Most patients experience only a mild and transitory illness, but severe neurologic complications have been described (Loos et al. 2014). ZIKV is predominantly transmitted by *Aedes* sp. mosquito species such as *A. aegypti* and *A. albopictus*, which also are vectors of dengue virus (DENV) and chikungunya virus (CHIKV). Laboratory diagnosis of ZIKV infection is based on IgG/IgM ELISA, and detection of viral RNA in the blood, serum or other body fluids (amniotic fluid, saliva, urine, semen), and/or viral isolation (Bachiller-Luque et al. 2016; Duffy et al. 2009). However, because of the co-circulation of arboviruses such as DENV and CHIKV, and cross-reactive antibodies, ZIKV infections are sometime missed or incorrectly diagnosed (Oehler et al. 2014).

In 2007, the first Zika epidemic occurred on the Yap Islands, an island chain in the Federated States of Micronesia. In 2013, a new epidemic started in French Polynesia (Musso et al. 2014), when it was observed a 20-fol increase in the incidence of Guillain-Barré syndrome (GBS), an immune mediated ascending flaccid paralysis (Oehler et al. 2014). In Brazil, ZIKV was first identified in 2015 in the State of Bahia in patients with clinical symptoms similar to those of dengue fever (Campos et al. 2015; Pinto Junior et al. 2015). These initial cases in Bahia were followed by epidemics in other northeastern states of Brazil. In the second half of 2015, there was a significant increase of newborns with microcephaly and other birth abnormalities in Northeastern Brazil, with 2,366 cases of microcephaly and other birth defects being reported by December of 2016, particularly in the State of Pernambuco (Schuler-Faccini et al. 2016). In Latin America, 1.6 million cases of ZIKV infections occurred between 2015 and 2016, followed by thousands of cases of microcephaly (WHO 2017; Oliveira et al. 2017).

The dramatic increase in the incidence of microcephaly concomitant with the ZIKV epidemic in Brazil alerted the scientific community to a potential causal association, leading the Brazilian Ministry of Health to declare the ZIKV epidemic to be a “national public health emergency”; and then, the World Health Organization (WHO) to declarate it to be a “public health emergency of international concern” (Albuquerque et al. 2018). Retrospective observational studies reinforced the suspicion of a causal of ZIKV infection with microcephaly (Araújo et al. 2018; Brasil et al. 2016; Cauchemez et al. 2016), and with non-congenital neuroinflammatory diseases, such GBS, encephalitis and myelitis (Cao-Lormeau et al. 2016; Parra et al. 2016; Styczynski et al. 2017; Salinas et al. 2017; Mehta et al. 2018). Several reports where instrumental in establishing the association between ZIKV and microcephaly. Mlakar et al. (2016) identified ZIKV in the brain tissue of a severe microcephalic stillbirth whose mother had been infected with ZIKV in Brazil. This report was one of first to propose the possibility of an association between ZIKV and microcephaly, which was later confirmed by case-control studies with Brazilian newborns (MERG Group 2016; Cordeiro et al. 2016; Brasil et al. 2016; Krow-Lucal et al. 2018).

Based on this literature, microcephaly combined with other fetal abnormalities associated with congenital ZIKV infection, in addition to microcephaly, have earned their own name: Congenital Zika Syndrome (CZS) (Miranda-Filho et al. 2016). Since then, CZS has been reported at variable frequencies in different investigations of pregnant women infected with ZIKV. Honein et al. (2017), studying 442 pregnancies with laboratory evidence of recent ZIKV infection in US territories, reported a 6% incidence of CZS including 18 cases of microcephaly (4%). In the case-control study by Hoen et al. (2018), in the French territories in the Americas, 7% of fetuses and infants of 527 ZIKV-PCR positive women were born with CZS, among which 32 (6%) had microcephaly. In turn, Brasil et al. (2016) had reported a very high frequency of 42% of CZS in infants born to 117 ZIKV-PCR positive woman from Rio de Janeiro in Southeastern Brazil, among which there were four babies with microcephaly (3.4%). It is noteworthy that this Brazilian case-control study applied an expanded CZS criterion, including children with tenuous and unspecific findings on MRI scans (Brasil et al. 2016).

These variations may be related, in part, to differences in the CZS/microcephaly diagnostic criteria, but variation in CZS risk could also explain these disparities. Furthermore, striking differences in incidence of microcephaly were observed in different regions of Brazil, the country with the most extensive ZIKV epidemic. More than 75% of the Brazilian cases of microcephaly were from the Northeast region, where the monthly incidence reached many times the incidence of the Southern population: 50 versus 1.5 per 10,000 live births (Oliveira et al. 2017). Accordingly, Barbeito-Andrés et al. (2018) suggested that cofactors, such as ZIKV strains, and environmental and human genetic variations, could potentially explain the geographic clustering of microcephaly in the Northeast of Brazil. They further showed an association between the incidence of microcephaly and the states with the highest prevalence of undernutrition (Barbeito-Andrés et al. 2020).

Studies in mice identified highly susceptible as well as resistant mouse strains as relevant models to access the mechanisms of ZIKV infection. Manet et al. (2020) showed that susceptibility was driven by multiple loci with small individual effects, and was independent of the Oas1b gene, which functions in the innate antiviral immunity pathway. Another recent animal study showed differences in inoculation routes influencing the rate of infection and viral load in mouse embryos (Duggal et al. 2018). These models reflect the role of host diversity on ZIKV infection outcome and highlight the need for additional studies in humans to address the role of different cofactors involved in CZS (Barbeito-Andrés et al. 2018). Genetic data related to human CZS are rare, in part due to difficulties in developing well-controlled case-control studies. Nonetheless, one revealing study found that discordant CZS twins have differential in vitro viral susceptibility of neural progenitor cells, expressing *MTOR* and *WNT* regulatory genes differently; both genes are part of key pathways in programming neural embryology (Caires-Junior et al. 2018). Case-control studies have shown the influence of maternal adenocyclase (*ADCY*) SNVs (Rossi et al. 2019) and of *TOL3* (maternal)/*TNFα* (infants) SNVs (Santos et al. 2019) in susceptibility of CZS and/or microcephaly. In turn, Azamor et al. (2018) indicated that a Type III interferon maternal SNV promotes protection against the development of CZS. Taken together, these data indicate that host genetic background related to antiviral responses, especially against arboviruses, may affect the outcome of fetal ZIKV infection.

Human genetic risk factors could be key elements in ZIKV-mother and ZIKV-baby interactions, as they might influence the vertical transmission of the virus, cell and tissue damage, and consequent congenital abnormalities associated with the ZIKV infection. However, the frequency of relevant alleles may not be uniformly distributed across global populations. In fact, population genetic differences likely arose during the evolution that occurred during the dispersion of *Homo sapiens* from Africa to other continents roughly 100,000 years ago. This dispersion led to selection of a series of genetic adaptations under different climatic and other environmental conditions, including food sources and local pathogens, leading to population differentiation. This differentiation, in turn, led to continuous gradients of genomic variants (Serre and Paabo 2004), including gradients of various resistance/susceptibility alleles for infectious diseases (Karlsson et al. 2014). We hypothesize that alleles at a number of loci in genes that influence immunity, interacting in a complex manner, may drive differences in the balance between resistance and susceptibility to CZS, forming a geographic genomic gradient of host susceptibility.

To test our hypothesis, we applied a methodologic approach that is pioneering in both method and concept. We developed a new tool based on SNV allele frequencies in several genes involved in immune function, evaluated together in a complex manner by machine learning techniques. Based on data from previous outbreaks of ZIKV, we hypothesized that sub-Saharan Africans were resistant to CZS and that Latin Americans were susceptible, these are the two populations that represent the theoretical extremes for risk of CZS epidemics. Our dual decision tree (DT)/trained machine learning models were able to distinguish individuals from Africa and the Americas with an accuracy of 98% using only ten SNVs from ten genes involved in immune functions (*CLEC4M* / *CD299, CD58, OAS2, CD80, VEPH1, CTLA4, CD274, CD209, PLAAT4, CREB3L1*). Here, we present the *Human Immune Genome Complex Gradient Theory* as potential explanation of why the CZS catastrophe occurred in Brazil.

## RESULTS

We developed five dual DT/machine learning classification pipelines (e.g. DT/SVM linear) exclusively with immune SNVs in genes involved in immune functions, aiming to detect a potential allele frequency gradient across sub-Saharan African, East Asian, and Latin American populations, and to determine if it was consistent with the CZS risks in ZIKV-infected women among these continents. The analysis of the SNVs (hypothetical candidates features) resulted in a set of ten SNVs from ten genes involved in immune functions: *CLEC4M/CD299, CD58, OAS2, CD80, VEPH1, CTLA4, CD274, CD209, PLAAT4*, and *CREB3L1*, which together displayed high power to discriminate between sub-Saharan Africans and Latin Americans. This selected SNV set provided high accuracy with all five dual DT/ML techniques (Fig. 1). The confusion matrices, ROC Curves, and other metrics suggest that this immune response SNV set is indeed a reliable discriminator of the two populations, which are theoretically at the extremes of risk for CZS.

**Fig. 1.**
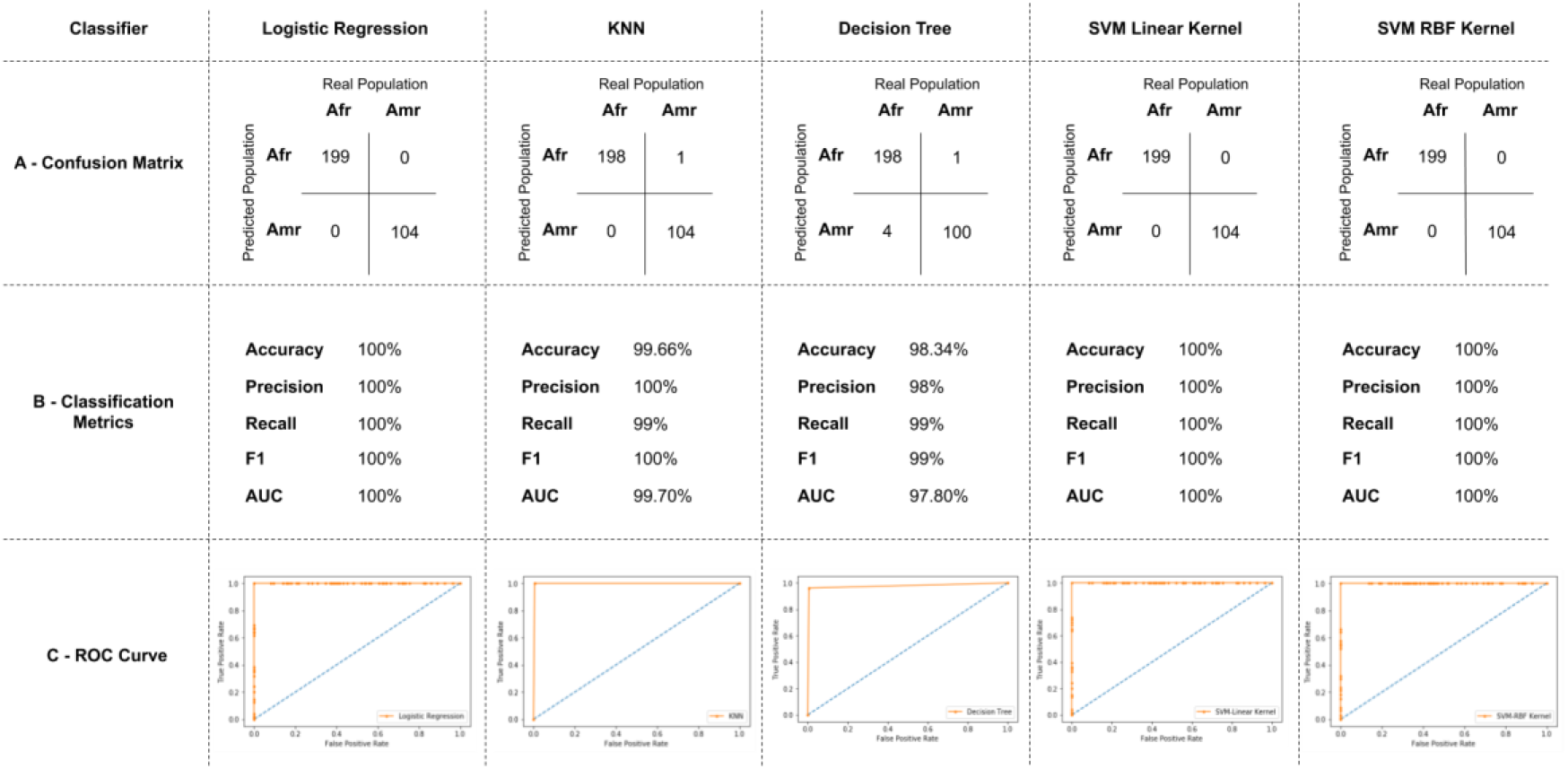
Statistical metrics (confusion matrix, classification metrics, ROC curve) for all five tested dual DT/ML genome approaches for immune genome discrimination between sub-Saharan African and Latin American populations: DT/Linear regression, DT/KNN, DT/DT, DT/SVM Linear, DT/SVM Kernel.

Figure 2 displays our *Human Immune Genome Complex Gradient Theory* as an infographic. Briefly, our dual DT/trained machine learning models, using the immune genome complex approach, revealed a unique and robust tool comprising ten SNVs acting in an integrated manner to discriminate sub-Saharan African and Latin American populations (Figure 2-A). For six of the ten SNVs, *VEPH1/rs3911403-T, CREB3L1/rs7127254-C, PLAAT4/rs4963271-A, CD58*/rs1335531-G, *CD80*/rs2629396-G, and OAS2/rs15895-A, the reference allele increased in frequency from sub-Saharan Africa to Latin America (Figure 2-B). In contrast, *CLEC4M*/rs12610506-G, *CD274*/rs4143815-G, and *CD209*/rs4804801-T decreased in frequency from sub-Saharan Africa to Latin America, while *CTLA4*/rs41265961-G was virtually constant. In addition to differences in host genetic variation at these ten SNPs, the circulating ZIKV lineage differed by continent (Figure 2-C) (Liu et al. 2019). Similarly, risk estimates of microcephaly in infants born to women infected with ZIKV during pregnancy differed by continent (Figure 2-D) (Cauchemez et al. 2016; Oliveira et al. 2017).

**FIG. 2.**
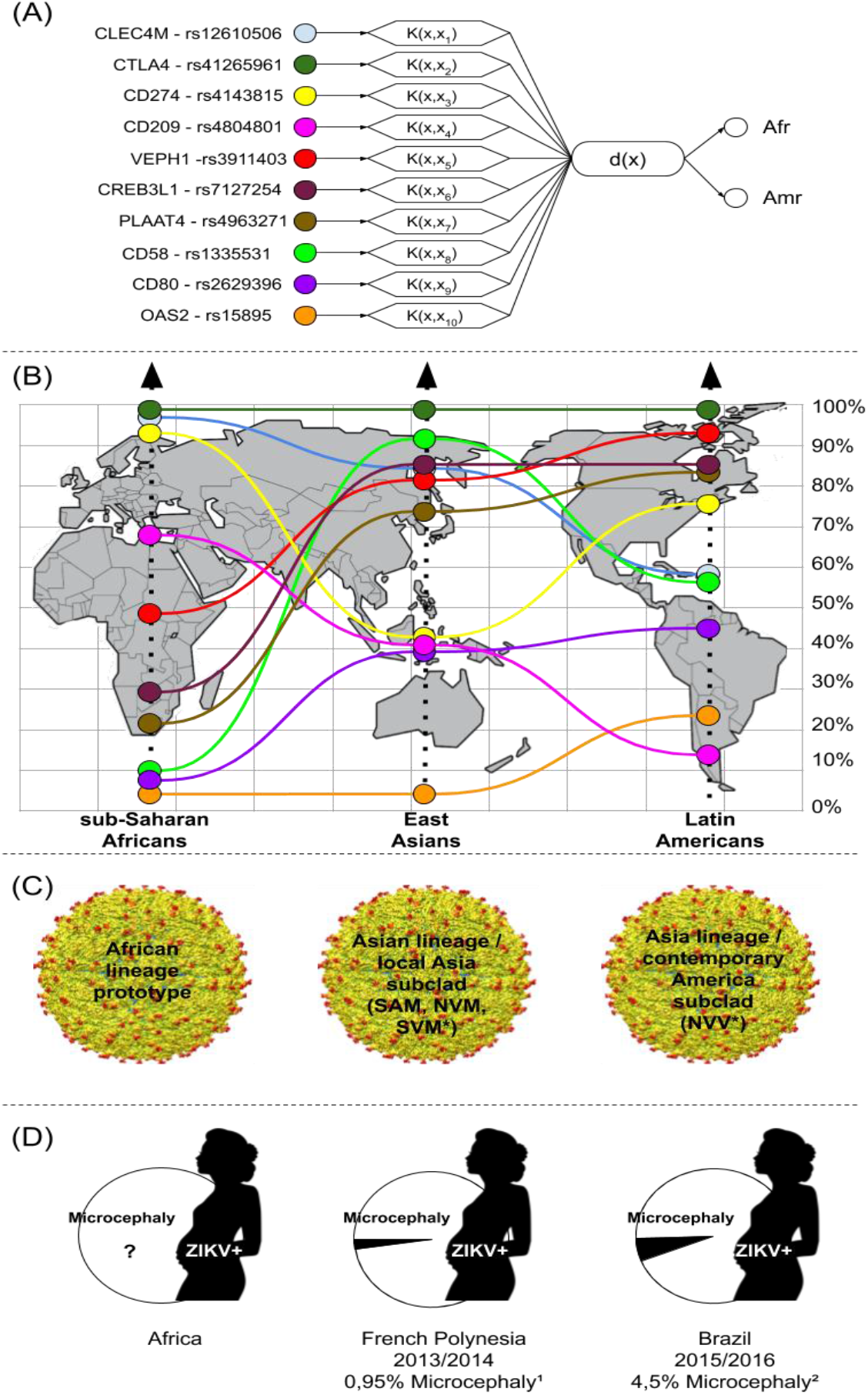
Machine learning approach and genetic CZS/microcephaly hypothesis overview. A) Genotype based Support Vector Machine (SVM) approach (second round) used for sub-Saharan African/Latin American population discrimination using only immune gene data pre-selected by DT, with ten SNVs and the hyperplane separation stage. B) Reference allele frequencies of the ten selected SNV/gene candidates for CZS phenotype predisposition in South America, showing allele frequency differences between sub-Saharan Africans and Latin Americans. C) ZIKV lineage circulating in each continent (*Asian Lineages are classified according to amino acid pattern at three positions: 139 in prM, 982 in NS1 and 2634 in NS5) (Liu et al. 2019). D) Probability of microcephaly in babies born to ZIKV-infected mothers during epidemics on the three continents (^1^Cauchemez et al. 2016; ^2^Oliveira et al. 2017).

Allele frequencies of the ten selected SNVs are shown for each 1000 Genomes continental (Table 1).

**Table 1.** Allele frequencies of ten selected SNVs in the DT/SVM complex immune genome tool among subSaharan African, East Asian, and Latin American populations from the 1000 Genomes.

| rsID | Gene | Chr:Position<br>(HG38) | Alleles<sup>a</sup> | sub-Saharan<br>Africans<br>(661) | East<br>Asians<br>(347) | Latin<br>Americans<br>(489) |
| --- | --- | --- | --- | --- | --- | --- |
| rs1335531 | *CD58* | 1:116572755 | G/C | 0.10 | 0.92 | 0.76 |
| rs41265961 | *CTLA4* | 2:203871138 | G/A | 0.99 | 1.00 | 0.99 |
| rs2629396 | *CD80* | 3:119530120 | G/T | 0.06 | 0.37 | 0.43 |
| rs3911403 | *VEPH1* | 3:157444338 | T/A | 0.47 | 0.80 | 0.92 |
| rs4143815 | *CD274* | 9:5468257 | G/C | 0.98 | 0.42 | 0.57 |
| rs7127254 | *CREB3L1* | 11:46286998 | C/T | 0.29 | 0.85 | 0.83 |
| rs4963271 | *PLAAT4* | 11:63537712 | A/G | 0.23 | 0.72 | 0.83 |
| rs15895 | *OAS2* | 12:113010483 | A/G | 0.01 | 0.00 | 0.22 |
| rs4804801 | *CD209* | 19:7742596 | T/A | 0.67 | 0.40 | 0.11 |
| rs12610506 | *CLEC4M/CD299* | 19:7768185 | G/A | 0.98 | 0.84 | 0.58 |

<sup>a</sup>The reference allele is listed first followed by the non-reference allele. All population allele frequencies are provided for the reference allele

## DISCUSSION

### CZS epidemiology

The CZS epidemic is recent and peaked in an epidemiological crisis in Brazil in 2015, becoming a more widespread geographic problem in 2016. Several reports were important to establish the causal link between ZIKV and microcephaly and other congenital malformations, which together became known as CZS. In recent CZS literature, there have been significant advances in our understanding of ZIKV epidemiology and evolution. This report presents a hypothesis to explain the high level of ZIKV teratovirulence and the high incidence of CZS cases in South America, especially in Brazil.

As of January 2017, 2,635 cases of central nervous system malformation potentially associated with ZIKV had been reported around the world, among which 2,366 were from Brazil and 78 were from Colombia (WHO 2017). Furthermore, although ZIKV might be widespread across Africa, reports of CZS cases in Africa are rare (WHO 2017; Hill et al. 2019). This dichotomy is not likely to be the consequence of differences in rate of ZIKV infection among pregnant women from different epidemic regions. A meta-analysis by Coelho et al. (2017), including observational studies, found a high inter-study variability in the frequencies of microcephaly in infants born to ZIKV-infected women, ranging from zero to 14%. This variability may be due to differences in the case definition of ZIKV infection and CZS between studies. However, some population-based estimates indicate that the risk of microcephaly may be different among ZIKV-infected pregnant women from different populations (Pacheco et al. 2016; Cauchemez et al. 2016; Oliveira et al. 2017; Barbeito-Andrés et al. 2020).

During the ZIKV outbreak in French Polynesia (2013-2014), only eight cases of microcephaly occurred, resulting in a risk of 0.95% among women infected in the first trimester of pregnancy (Cauchemez et al. 2016). Impressively, 1,950 cases of infection-related microcephaly were confirmed after ZIKV epidemics in Brazil (2015-2016), reaching an incidence of 4.5% in babies born to infected woman (Oliveira et al. 2017). It is important to highlight that, in the Brazilian epidemics (2015-2016), 31,000 ZIKV suspected cases were reported to the surveillance systems, but it is estimated that 1,673,272 persons were infected with ZIKV (Oliveira et al. 2017). Similarly, in the Polynesian epidemic 31,000 persons were estimated to have been infected, but only 8,000 cases of ZIKV infections were officially reported (Cauchemez et al. 2016). Although we cannot account in our analysis for the under-reporting in our consideration of the incidence of CZS, there were strikingly different risks even within Brazil, the country with the most extensive ZIKV epidemics. More than 75% of the Brazilian cases of microcephaly were from the Northeast region, where the monthly incidence reached several times the incidence of Southern population: 50 *versus* 1.5 per 10,000 live births (Oliveira et al. 2017). In turn, an ecological study at a municipality level across Brazil was unable to correlate the incidence of microcephaly with the incidence of ZIKV infection, but this showed a correlation with poverty index. These data suggested that ZIKV might not be, alone, be responsible for the unusual pattern of microcephaly in Brazil that clustered in the Northeast region (Campos et al. 2018).

Some hypotheses have emerged to explain the asymmetrical distribution of CZS risk and ZIKV neurovirulence. The evolution of ZIKV strains, vector competence of mosquitoes across continents, climatic conditions, environmental toxins, and nutritional factors deserve to be considered. In turn, immunological and genetic variability influencing the susceptibility/resistance status of the human host is also worthy of concern, especially in ZIKV naïve areas where DENV has been hyper-endemic, as pre-infection with DENV might lead to an antibody enhancement response that induces the severest clinical outcomes (Zhong-Yu et al. 2019; Barbeito-Andrés et al. 2020). To date, there is a single ZIKV serotype, but two major viral lineages (African and Asian) have been identified through phylogenetic analyses. The Asian lineage can be divided into two subclades (Asian and Contemporary American), which are responsible for the recent outbreaks and are associated with clinical severity in the Pacific region and the Americas, respectively (Liu et al. 2019). Some experimental studies based on reverse genetics have shown that ZIKV undergoes genetic changes over time and that amino acid mutation patterns in ZIKV surface proteins may have contributed to the epidemic waves that occurred in Micronesia (2007), French Polynesia (2013-2014), and South America (2015-2016) (Faria et al. 2017; Metsky et al. 2017). Among the genetic changes, three important amino acid mutations, respectively, in prM S139N (substitution of Serine to Asparagine at position 139), NS1 A928V (substitution of Alanine to Valine), and NS5 M/T2634V (substitution of Methionine or Threonine to Valine), have been related to a more neurovirulent and pathogenic ZIKV strain spreading from Asia to the Americas (Zhong-Yu et al. 2019) (Fig. 1). An increase in neurovirulence and pathogenic capacity of ZIKV as it spread to the Americas may have been a contributing factor, together with the host genetic factors that we propose, to the CZS catastrophe in Brazil.

Among environmental cofactors, contaminants in the water supplies, pre-natal vaccines, and the use of alcohol, cigarettes, or drugs during pregnancy were all considered as potential contributing factors to the Brazilian cluster of microcephaly at the beginning of epidemic. In the first case-control study proving the association of microcephaly with ZIKV in Pernambuco (OR= 73; CI 13-∞), neither vaccination during pregnancy or use of drugs, alcohol, or larvicides, were associated with microcephaly (Araújo et al. 2018). However, smoking was found to be associated with microcephaly (Araújo et al. 2018). Similarly, an ecological study found no correlation between the use of the larvicide pyriproxyfen in stored water and microcephaly among municipalities of the metropolitan region of Pernambuco, the epicenter of the Brazilian microcephaly epidemic (Albuquerque et al. 2016). Interestingly, experiments in mouse models suggested that maternal diet protein restriction (Barbeito-Andrés et al. 2020) and exposure to cyanobacterial saxitoxin through water intake (Pedrosa et al. 2019) exacerbate brain malformations induced by ZIKV. In fact, CZS may have multiple environmental cofactors and may be influenced by ZIKV genetic factors, but the constellation of alleles of host genes involved in immune functions may also be important predictor of the occurrence and severity of CZS.

### CZS genetics and the hypothetical *Human Immune Genome Complex Gradient*

Genetic data pertinent to CZS are rare due the difficulties in recruiting human CZS cohorts into well-controlled CZS case-control studies. Developing such studies is particularly challenging given the difficulties associated with diagnostic certainty of congenital ZIKV-infection as well as ascertaining the embryonic or fetal period of infection. In fact, not a single well-controlled genomic study was conducted with the hundreds of children with CZS during the Brazilian epidemics. However, genetic studies using animal models can shed light on the role of different cofactors involved in CZS. IFN I receptor gene (*IfnarI*) knockout mice and the use of anti-IFNAR antibody *in vivo* showed that CZS susceptibility was driven by multiple loci with small individual effects and independent of the *Oas1b* gene (Manet et al. 2020). These results indicate that novel experimental models of infection can help to address the role of different cofactors and genes involved in CZS.

Given the paucity of genetic data from human ZIKV case-control association studies, and combined with evidence from animal models demonstrating the impact of host genetic factors on the development of CZS, we are presenting a new theory to help explain the Brazilian CZS catastrophe of 2015-2016: the *Human Immune Genome Complex Gradient* hypothesis. Our analysis of global allele frequency data from 297 immune response genes across three continental populations suggest that, at the population level, sub-Saharan Africans are quite resistant to CZS, East Asians are moderately susceptible, and Latin Americans are highly susceptible to CZS. The rationale is based on ten key SNPs in immune functional genes, whose alleles collectively discriminate among three populations, creating a combined signature for each, and forming a hypothetical geographical gradient for risk of CZS. The genes exhibiting these SNPs (*CLEC4M* / *CD299, CD58, OAS2, CD80, VEPH1, CTLA4, CD274, CD209, PLAAT4, CREB3L1*) are important for function of innate immune system and for anti-viral response. We posit that, in a complex manner, the constellation of SNVs in these genes can potentially influence ZIKV vertical transmission, tissue damage, and congenital abnormalities; and consequently, the outcome of ZIKV-mother and ZIKV-fetus interactions.

We advocate that an immune genome gradient is probably developed as consequence of host-pathogen co-evolution during *H. sapiens* dispersion across continents thousands of years ago. Modern humans appeared in Africa about 200,000 years ago, dispersed to Asia approximately 100,000 years ago, and subsequently colonized the rest of the world in a series of migratory events. During these events, humans encountered a wide range of different environmental conditions, such as climate, food availability, and local pathogens (Fumagalli et al. 2011). These environmental differences probably selected for genetic adaptations and resulted in the establishment of continuous genome gradients of allele frequencies across populations (Serre and Paabo 2004; Marian 2009).

For infectious diseases, natural selection in human populations is evident for pathogens with a long-standing relationship with *H. sapiens*, including those that cause malaria, smallpox, cholera, tuberculosis, and leprosy (Karlsson et al. 2014). The timing, strength and direction of selection (that is, positive, negative, or balancing) shape the patterns of genomic variation (Karlsson et al. 2014). Hence, mutations in immune-inflammatory response genes are good candidates for understanding why some populations are more susceptible to infections than others. For example, in models of human genome adaptations, approaches that consider a single pathogen have provided examples of host evolution driven by the influence of that pathogen. The case of anemia *falciform* is a classic example. In heterozygosity, the sickle cell allele confers sickle cell trait and protection against *Falciparum malaria*, while in homozygotes it is life threatening; and so a sickle cell trait occurs at a high frequency in malaria-rich regions, such as Africa, and at a low frequency outside of these regions (Kwiatkowski 2005). Another inherited red blood cell disorder that is associated with malaria resistance is caused by glucose-6-phosphate dehydrogenase (G6PD) deficiency, which also coincides with the geographical distribution of malaria (Tishkoff et al. 2001).

Many other instances of population level differences in infectious disease susceptibility have been described. For example, MHC complex (HLA) diversity confers differential resistance to a wide range of pathogens (Buhler and Sanchez-Mazas 2011), and has been demonstrated to impact the clinical course of infection for a large number of viral infectious diseases including dengue fever (Stephens et al. 2010), yellow fever (Co et al. 2002), and hepatitis C Virus (HCV) (Kuniholm et al. 2011). Genetic variation in toll like receptor genes suggests that Europeans exhibit a reduced TLR1-mediated response compared to other global populations (Barreiro et al. 2009). Another example, African Americans are more likely than Caucasians to carry cytokine variants that up-regulate pro-inflammatory responses (Ness et al. 2004). This may predispose individuals of African ancestry to higher rates of inflammatory diseases (Ness et al. 2004). Similarly, Pennington et al. (2009) suggested that African populations from tropical Africa have a more robust inflammatory response compared to Europeans and East Asians. They hypothesized that this robust response was the result of variation in immune response genes that evolved in response to local pathogen exposure: parasites with low virulence and long lifespans such as those that cause malaria in Africa versus high-virulence pathogens such as those that cause smallpox and cause measles in Europe and East Asia. Taken together, these are emblematic examples demonstrating that human genetic variation contributes to differences in response to infections, and they support our hypothesis that a complex genome gradient has contributed to CZS phenotype distribution across global populations.

There have been studies that examined the impact of immune genetic factors impact on CZS. Hamel et al. (2015) carried out an *in vitro* gene expression study on ZIKV infection in epidermal cells, and demonstrated that receptors such as DC-SIGN, TIM-1, TIM-4, AXL, and TIROL are important elements for ZIKV entry into cells. They observed significant reduction in the percentage of ZIKV-infected cells when the AXL receptor is blocked by antibodies, and similarly for DENV-infected cells. The ZIKV-infected fibroblasts presented expression of anti-viral response genes, cell receptors genes, and immunity genes. Up-regulation of many genes involved in immune response has also already been observed in models of DENV infection, such as *FAAD, IFN, MYD88* (Gomes et al. 2010), *OAS, PYCARD, TNF* (Nascimento et al. 2009), and *STAT* (Warke et al. 2003), supporting the idea of common key elements for both flaviviruses. In humans, case-control immune genome studies have shown the influence of maternal adenocyclase SNV (Rossi et al. 2018) and TOL3 (maternal)/TNFα (infants) (Santos et al. 2019) on CZS susceptibility or severe microcephaly. On the other hand, Azamor et al. (2018) indicated that a maternal IFN III SNV promotes protection against the development of CZS. However, to date, there are no reports regarding influence of a combination of genotypes at multiple loci/genes on the risk for CZS.

### The immune genome SNV tool

Our dual DT/ML immune genome approach showed that allele frequencies of ten SNPs of ten genes are consistent with the risk of microcephaly across Africa, French Polynesia, and Brazil, and may have contributed to the Brazilian CZS epidemics. All of these genes (*CLEC4M, CD58, OAS2, CD80, VEPH1, CTLA4, CD274, CD209, PLAAT4, CREB3L1*) are involved with innate immune response, virus cycle, or anti-virus response. Seven of those polymorphisms are genic SNPs (six are “transcript variants” and one is “stop codon lost”), rather than in non-genic sequence. Our results agree with Fumagalli et al. (2011) who found a significant enrichment of genic SNPs, in particular non-synonymous SNPs *vs* intergenic SNPs, involved in local adaptation to pathogenic exposure across human populations. As Karlsson et al. (2014) pointed out that much adaptive evolution probably occurred in regulatory, non-coding regions, but our results indicate that genic variations may be more frequent in the human adaptive response to pathogenic exposure. It is also important to point out that most studies identifying host genetic resistance factors to pathogenic infection show one or a few SNVs functioning in an individual manner. Our findings suggest that genetic variations in many genes may work in an integrated and coordinated way, thus indicating polygenic selection.

It is noteworthy to state that our DT/ML selected-SNPs comprise a broad functional panel. Type II C-type lectin (CLEC4C) is a transmembrane protein expressed selectively on plasmacytoid dendritic cells. The triggering of the extra-cellular C-terminal C-type carbohydrate recognition region of CLEC4C regulates the secretion of proinflammatory cytokines and type I interferons (IFNs) (Lim et al. 2016), directly related with the oligoadenylate synthetase (OAS) family. OAS proteins detect foreign RNA by oligoadenylate cytosolic PAMP (Pathogen Associated Molecular Pattern), which leads to 2-5-oligoadenylate synthesis, which then acts as an intracellular second messengers to activate latent ribonuclease L (RNase L). This activation promotes indiscriminate cleavage of both host and viral RNA, and the production of additional PAMPs, which reinforces the innate immune response (Schneider et al. 2014, Chakrabarti et al. (2011). Upon stimulation, Toll-Like Receptors (TLRs) 7, 8, and 9 interact with the TIR domain of MyD88 to recruit signaling molecules Interleukin-1 receptor-associated kinases (IRAK4/1/2) (He et al. 2013). This recruitment leads to IkB degradation and thereby contributes to the activation of NF-kB and MAPK signaling pathways to induce secretion of inflammatory cytokines, IFN type I, chemokines, and antimicrobial peptides, and other innate immune response genes (He et al. 2013). Some of these genes have been previously related with dengue fever, such as CLEC4M, OAS2,3, VEPH1 (Davi et al. 2019), CD209 (Tassaneetrithep et al. 2003), and Zika infection, such as CD80, TLR7 (Hamel et al. 2015).

It is important to consider that the individual humans are “complex systems” programed by billions of DNA base pairs. During evolution, DNA replication errors resulted in many mutations across generations, and some favorable mutations became retained in the genome as consequence of natural selection, contributing to population diversity. Here, our principal message regarding host influences on CZS, beyond the immune genome gradient itself, is the complexity of inter-individual and inter-population genetic backgrounds. We are proposing a new theory to help explain the CZS catastrophe in Brazil. The hypothetical “human immune genome complex gradient” is based on the host genetic population gene diversity produced by human natural history and evolution across the world. Going forward, we propose that the complexity of genetic data should be deeply analyzed using the most robust available computational tools and a much broader immune genome SNV panel. Furthermore, ZIKV genetic factors, environmental factors, and population immune history are essential in the modeling the complex clinical phenotype of CZS Brazilian phenomenon.

## METHODS

In order to examine the CZS phenomenon in Brazil from an evolutionary perspective, we created a theoretical population immune genome in silico analysis. This method used a dual computational approach: (i) a Decision Tree (DT) to select SNVs (features) as good population discriminants and (ii) other machine learning models to improve the discrimination quality of the selected SNV cluster. We assume theoretical (pooled, probably admixed) populations in this approach. We consider sub-Saharan Africans, who evolved (probably) together with ZIKV, to be naturally resistant; and we consider Latin Americans to be susceptible due to their historical ZIKV naïve condition and the recent epidemics of CZS. As such, these two populations represent the theoretical genetic extremes of CZS risk. In addition, we hypothesize that East Asians are moderately susceptible.

### Dataset characterization and preprocessing

The dataset was composed of genotype data from The 1000 Genomes Project (2015), and included individuals from a mix of populations from three continents: the Americas, Africa, and Asia (https://www.internationalgenome.org). The American population consisted of Mexicans living in Los Angeles, United States (MXL), Puerto Ricans from Puerto Rico (PUR), Colombians from Medellin (CLM) and Peruvians from Lima (PEL). The East Asian population was comprised of Kinh from Ho Chi Minh City, Vietnam (KHV), Han Chinese from South China (CHS), Han Chinese from Beijing (CHB), Chinese Dai from Xishuangbanna (CDX), and Japanese from Tokyo (JPT). The African population was composed of Esan (ESN) and Yoruba (YRI) from Nigeria, Luhya from Webuye, Kenya (LWK), Gambian Mandinka (GWD), Mende from Sierra Leone (MSL), and descendants of Africans living in Barbados (ACB) and in the southwestern United States (ASW). From each population, we selected 297 innate immune system SNVs previously interrogated in a dengue genetic study (Davi et al. 2019). In total, the dataset comprised 297 attributes, corresponding to 297 target SNVs and 1497 individuals: 661 Africans, 347 East Asians, and 489 Latin Americans. Dataset preprocessing was performed according to the following definitions: (i) Each SNV was evaluated considering both alleles as a single point of information; (ii) heterozygote information was homogenized (e.g. “A”/”C” and “C”/”A” were considered the same); (iii) missing data were assigned the value NN. All attributes were categorical, containing information about allelic pairs of each considered SNP - for example homozygotes such as AA, or heterozygotes such as AT. For each SNP, each pair of alleles received an integer code. For example, a given SNP containing allelic pairs AA, AT, and TT, was encoded as follows: AA = 1, AT = 2, and TT =3. This integer encoding was done independently for each SNP.

### Feature selection and classification

Decision Trees (DT) (Utgoff 1989) were used in the first step to reduce the prior set containing 297 SNVs to a smaller SNV dataset with a higher power for discrimination between sub-Saharan Africans and Latin Americans. This was done to take advantage of the DT characteristic to naturally select features that best discriminate classes, in this case distinct populations. In addition, DTs make it possible to quantify and rank the importance of each attribute, which is desirable for the subsequent biological analysis. Stratified 10-fold cross-validation was performed as part of the feature selection effort. In this process, the dataset was divided into ten parts of which nine were used as the training set and one part was used as the test set. This procedure was repeated ten times, so that each pattern appeared only once as part of the test set. Stratification occurred so that the proportion of individuals in each class in the training and test sets was similar to the proportion of the dataset as a whole. After completing the ten repetitions, the SNVs that were used by DT analysis in at least six of the ten repetitions were considered as potentially good population discriminators and were selected to be used in the second step by other classifier algorithms.

As the second step, the selected SNVs were evaluated regarding their discrimination quality. Algorithms with wide acceptance in the literature were used: (i) logistic regression (LR) (Cox 1966), (ii) method of k nearest neighbors (KNN) (Cover 1967) and (iii) support vector machines (Boser et al. 1992) with linear kernel (SVM-Linear) and with radial base function kernel (SVM-RBF). Hence, we employed five dual DT/machine learning classification pipelines: DT/LR, DT/KNN, DT/DT, DT/SVM-Linear, DT/SVM-RBF. The selected SNV set was evaluated using holdout technique; accuracy and precision were calculated, and confusion matrixes and ROC-Curves (correlation curve between Sensibility and 1-Specificity) for each model were generated.

## ACKNOWLEDGMENTS

We would like to thank the financial support of the FIOCRUZ/CNPQ PAPES VII program (grant no.41910/2015-6).

## REFERENCES

Albuquerque MFPM, Souza WV, Araújo TVB, Braga MC, Miranda-Filho DB, Ximenes RAA, Melo-Filho DA, Brito CAA, Valongueiro S, Melo APL, et al. 2018. The microcephaly epidemic and Zika virus: building knowledge in epidemiology. Cad Saúde Pública 2018 34: e00069018. doi: 10.1590/0102-311X00069018

Albuquerque MFPM, Souza WV, Mendes ACG, Lyra TM, Ximenes RAA, Araújo TVB, Braga C, Miranda-Filho DB, Martelli CMT, Rodrigues LC. 2016. Pyriproxyfen and the microcephaly epidemic in Brazil - an ecological approach to explore the hypothesis of their association. Mem Inst Oswaldo Cruz 111: 774–776.

Araújo TVB, Ximenes RAA, Miranda-Filho DB, Souza WV, Montarroyos UR, Melo APL, Valongueiro S, Albuquerque MFPM, Braga C, Brandão-Filho SP, et al. 2018. Association between microcephaly, Zika virus infection, and other risk factors in Brazil: final report of a case-control study. Lancet Infect Dis 18: 328–336. doi: 10.1016/S1473-3099(17)30727-2

Azamor T, Cunha DP, Silva AMV, Bezerra OCL, Ribeiro-Alves M, Calvo TL, Kehdy FSG, Manta FSN, Pinto TG, Ferreira LP, et al. 2019. Congenital Zika Syndrome is associated with maternal genetic background. bioRxiv doi: 10.1101/715862.

Bachiller-Luque, P., et al., First case of imported Zika virus infection in Spain. Enferm Infecc Microbiol Clin, 2016. 34(4): p. 243–6.

Barbeito-Andrés J, Pezzuto P, Higa LM, Dias AA, Vasconcelos JM, Santos TMP, et al. 2020. Congenital Zika syndrome is associated with maternal protein malnutrition. Sci Adv 6: eaaw6284. doi: 10.1126/sciadv.aaw6284

Barbeito-Andrés J, Schuler-Faccini L, Garcez PP. 2018. Why is congenital Zika syndrome asymmetrically distributed among human populations? PLoS Biol 16: e2006592. doi: 10.1371/journal.pbio.2006592

Barreiro L.B, Ben-Ali M, Quach H, Laval G, Patin E, Pickrell JK, Bouchier C, Tichit M, Neyrolles O, Gicquel B, et al. 2009. Evolutionary dynamics of human Toll-like receptors and their different contributions to host defense. PLoS Genet 5: e1000562. doi: 10.1371/journal.pgen.1000562

Boser BE, Guyon IL, Vapnik VN. 1992. A training algorithm for optimal margin classifiers. Proceedings of COLT ‘ 92: 144–152. doi: 10.1145/130385.130401

Brasil P, Pereira Jr JP, Moreira ME, Ribeiro Nogueira RM, Damasceno L, Wakimoto M, Rabello RS, Valderramos SG, Halai UA, Salles TS, et al. 2016. Zika virus infection in pregnant women in Rio de Janeiro. N Engl J Med 375: 2321–2334. doi: 10.1056/NEJMoa1602412

Buhler S and Sanchez-Mazas A. 2011. HLA DNA sequence variation among human populations: molecular signatures of demographic and selective events. PLoS One 6: e14643. doi: 10.1371/journal.pone.0014643

Caires-Júnior LC, Goulart E, Melo US, Araujo BHS, Alvizi L, Soares-Schanoski A, Oliveira DF, Kobayashi GS, Griesi-Oliveira K, Musso CM, et al. 2018. Discordant congenital zika syndrome twins show differential in vitro viral susceptibility of neural progenitor cells. Nat Commun 9: 475. doi: 10.1038/s41467-017-02790-9

Campos GS, Bandeira AC, Sardi SI. 2015. Zika virus outbreak, Bahia, Brazil. Emerg Infect Dis 21: 1885–1886. doi: 10.3201/eid2110.150847

Campos MC, Dombrowski JG, Phelan J, Marinho CRF, Hibberd M, Clark TG, Campino S. 2018. Zika might not be acting alone: Using an ecological study approach to investigate potential co-acting risk factors for an unusual pattern of microcephaly in Brazil. PLoS One 13: e0201452. doi: 10.1371/journal.pone.0201452

Cao-Lormeau VM, Blake A, Mons S, Lastère S, Roche C, Vanhomwegen J, Dub T, Baudouin L, Teissier A, Larre P, Vial A. et al. 2016. Guillain-Barre Syndrome outbreak associated with Zika virus infection in French Polynesia: a case-control study. Lancet 387:1531–1539. doi: 10.1016/S0140-6736(16)00562-6

Cauchemez S, Besnard M, Bompard P, Dub T, Guillemette-Artur P, Eyrolle-Guignot D, Salje H, Van Kerkhove MD, Abadie V, Garel C, Fontanet A. et al. 2016. Association between Zika virus and microcephaly in French Polynesia, 2013-15: a retrospective study. Lancet. 387: 2125–2132. doi: 10.1016/S0140-6736(16)00651-6

Chakrabarti A, Jha BK, Silverman RH. 2011. New insights into the role of RNase L in innate immunity, J. Interferon Cytokine Res 31: 49–57. doi: 10.1089/JIR.2010.0120

Co MD, Terajima M, Cruz J, Ennis F, Rothman A. 2002. Human cytotoxic T lymphocyte responses to live attenuated 17D yellow fever vaccine: identification of HLA-B35-restricted CTL epitopes on nonstructural proteins NS1, NS2b, NS3, and the structural protein E. Virology 293: 151–163. doi: 10.1006/VIRO.2001.1255

Coelho AVC and Crovella S. 2017. Microcephaly prevalence in infants born to zika virus-infected women: A systematic review and meta-analysis. Int J Mol Sci 18: 1714. doi: 10.3390/ijms18081714

Cordeiro MT, Pena LJ, Brito CA, Gil LH, Marques ET. 2016. Positive IGM for zika virus in the cerebrospinal fluid of 30 neonates with microcephaly in Brazil. Lancet 387: 1811–1812. doi: 10.1016/S0140-6736(16)30253-7

Cover TM and Hart PE. 1967. Nearest neighbor pattern classification. IEEE Trans Inf Theory 13: 21–27. doi: 10.1109/TIT.1967.1053964

Cox DR. 1966. Some procedures connected with the logistic qualitative response curve. Research Papers in Probability and Statistics (Festschrift for J. Neyman). (David FN ed.) 55–71. Wiley, London.

Davi CCM, Pastor A, Oliveira T, Lima Neto FB, Braga-Neto UM, Bigham A, Bamshad M, Marques ETA, Acioli-Santos B. 2019. Dengue Severity Prognosis Using Human Genome Data and Machine Learning Techniques. IEEE Trans Biomed Eng doi: 10.1109/TBME.2019.2897285.

Dick GWA. 1953. Epidemiological notes on some viruses isolated in Uganda (Yellow fever, Rift Valley fever, Bwamba fever, West Nile, Mengo, Semliki forest, Bunyamwera, Ntaya, Uganda S and Zika viruses. Trans R Soc Trop Med Hyg 47: 13–48. doi: 10.1016/0035-9203(53)90021-2

Dick GW, Kitchen SF, Haddow AJ. 1952. Zika Virus (I). Isolations and serological specificity. Trans R Soc Trop Med Hyg 46: 509–520. doi: 10.1016/0035-9203(52)90042-4

Duffy MR, Chen T, Hancock T, Powers AM, Kool JL, Lanciotti RS, Pretrick M, Marfel M, Holzbauer S, Dubray C, et al. 2009. Zika virus outbreak on Yap Island, Federated States of Micronesia. N Engl J Med 360: 2536-2543. doi: 10.1056/NEJMoa0805715

Duggal NK, McDonald EM, Ritter JM, Brault AC. 2018. Sexual transmission of Zika virus enhances in utero transmission in a mouse model. Sci Rep 8: 1–8. doi: 10.1038/s41598-018-22840-6

Faria NR, Quick J, Claro IM, Thézé J, de Jesus JG, Giovanetti M, Kraemer MUG, Hill SC, Black A, da Costa AC, et al. 2017. Establishment and cryptic transmission of Zika virus in Brazil and the Americas. Nature 546: 406–410. doi: 10.1038/nature22401

Fumagalli M, Sironi M, Pozzoli U, Ferrer-Admettla A, Pattini L, Nielsen R. 2011. Signatures of environmental genetic adaptation pinpoint pathogens as the main selective pressure through human evolution. PLoS Genet 7: e1002355. doi: 10.1371/journal.pgen.1002355

Gomes ALV, Wee LJK, Khan AM, Gil LHVG, Marques ETA, Calzavara-Silva CE, Tan TW. 2010. Classification of dengue fever patients based on gene expression data using support vector machines. PLoS One 5: e11267. doi: 10.1371/journal.pone.0011267

Hamel R, Dejarnac O, Wichit S, Ekchariyawat P. Neyret A, Luplertlop N, Perera-Lecoin M, Surasombatpattana P, Talignani L, Thomas F, et al. 2015. Biology of Zika virus infection in human skin cells. J Virol 89: 8880–8896. doi: 10.1128/JVI.00354-15

He X, Jia H, Jing Z, Liu D. 2013. Recognition of pathogen-associated nucleic acids by endosomal nucleic acid-sensing toll-like receptors. Acta Biochim Biophys Sin 45: 241–258. doi: 10.1093/abbs/gms122

Hill SC, Vasconcelos J, Neto Z, Jandondo D, Zé-Zé L, Aguiar RS, Xavier J, Thézé J, Mirandela M, Cândido ALM, et al. Emergence of the Asian lineage of Zika virus in Angola: an outbreak investigation. Lancet Infect Dis 2019 19: 1138–1147. doi: 10.1016/S1473-3099(19)30293-2

Hoen B, Schaub B, Funk AL, Ardillon V, Boullard M, Cabié A, Callier C, Carles G, Cassadou S, Césaire R, et al. 2018. Pregnancy outcomes after ZIKV infection in French Territories in the Americas. N Engl J Med 378: 985–994. doi: 10.1056/NEJMoa1709481

Hoffman SC, Stanley EM, Cox ED, DiMercurio BS, Koziol DE, Harlan DM, Kirk AD, Blair PJ. 2002. Ethnicity greatly influences cytokine gene polymorphism distribution. Am J Transplant. 2: 560–567. doi: 10.1034/j.1600-6143.2002.20611.x

Honein MA, Dawson AL, Petersen EE, Jones AM, Lee EH, Yazdy MM, Ahmad N, Macdonald J, Evert N, Bingham A, et al. 2017. Birth defects among fetuses and infants of US women with evidence of possible zika virus infection during pregnancy. JAMA 317: 59–68. doi: 10.1001/jama.2016.19006

Ioos S, Mallet H-P, Goffart IL, Gauthier V, Cardoso T, Herida M. 2014. Current Zika virus epidemiology and recent epidemics. Med Mal Infect 44: 302–307. doi: 10.1016/J.MEDMAL.2014.04.008

Karlsson EK, Kwiatkowski DP, Sabeti PC. 2014. Natural selection and infectious disease in human populations. Nat Rev Genet 15: 379–93.

Krow-Lucal ER, Andrade MR, Cananéa JNA, Moore CA, Leite PL, Biggerstaff BJ, Cabral CM, Itoh M, Percio J, Wada MY, et al. 2018. Association and birth prevalence of microcephaly attributable to Zika virus infection among infants in Paraíba, Brazil, in 2015–16: a case-control study. Lancet Child Adolesc Health 2: 205–213. doi: 10.1016/S2352-4642(18)30020-8

Kuniholm MH, Gao X, Xue X, Kovacs A, Marti D, Thio CL, Peters MG, Greenblatt RM, Goedert JJ, Cohen MH, et al. 2011. The relation of HLA genotype to hepatitis C viral load and markers of liver fibrosis in HIV-infected and HIV-uninfected women. J Infect Dis 203: 1807–1814. doi: 10.1093/INFDIS/JIR192

Kwiatkowski DP. 2005. How malaria has affected the human genome and what human genetics can teach us about malaria. Am J Hum Genet 77:171–192. doi:10.1086/432519

Lim SM, Kim YE, Choi WJ, Oh K-W, Noh M-Y, Kwon M-S, Nahm M, Kim N, Ki C-S, Kim SH, et al. 2016. CLEC4C p.K210del variant causes impaired cell surface transport in plasmacytoid dendritic cells of amyotrophic lateral sclerosis. Oncotarget 7: 24942–24949. doi:10.18632/oncotarget.7886

Liu Z, Shi W, Qin C. 2005. The evolution of Zika virus from Asia to the Americas. Nat Rev Microbiol 17: 131-139. doi: 10.1038/s41579-018-0134-9

Manet C, Simon-Lorière E, Jouvion G, Hardy D, Prot M, Conquet L, Flamand M, Panthier J-J, Sakuntabhai A, Montagutelli X. 2020. Genetic diversity of Collaborative Cross mice controls viral replication, clinical severity and brain pathology induced by Zika virus infection, independent of *Oas1b*. J Virology 94: e01034–19. doi: 10.1128/JVI.01034-19

Marian AJ. 2009. Nature’s genetic gradients and the clinical phenotype. Circ Cardiovasc Genet 2: 537–539. doi: 10.1161/CIRCGENETICS.109.921940

Mehta R, Soares CN, Medialdea-Carrera R, Ellul M, da Silva MTT, Rosala-Hallas A, Jardim MR, Burnside G, Pamplona L, Bhojak M, et al. 2018. The spectrum of neurological disease associated with Zika and chikungunya viruses in adults in Rio de Janeiro, Brazil: A case series. PLoS Negl Trop Dis 12: e0006212. doi: 10.1371/journal.pntd.0006212

MERG Group. 2016. Microcephaly in infants, Pernambuco State, Brazil, 2015. Emerg Infect Dis 22: 1090–1093. doi: 10.3201/EID2206.160062

Metsky HC, Matranga CB, Wohl S, Schaffner SF, Freije CA, Winnicki SM, West K, Qu J, Baniecki ML, Gladden-Young A, et al. 2017. Zika virus evolution and spread in the Americas. Nature 546: 411–415. doi: 10.1038/nature22402

Mlakar J, Korva M, Tul N, Popović M, Poljšak-Prijatelj M, Mraz J, Kolenc M, Rus KR, Vipotnik TV, Vodušek VF, et al. 2016. Zika virus associated with microcephaly. N Engl J Med 374: 951–958. doi: 10.1056/NEJMoa1600651

Musso D, Nilles EJ, Cao-Lormeau VM. 2014. Rapid spread of emerging Zika virus in the Pacific area. Clin Microbiol Infect 20: O595–6. doi: 10.1111/1469-0691.12707

Miranda-Filho DB, Martelli CMT, Ximenes RAA, Araújo TVB, Rocha MAW, Ramos RCF, Dhalia R, França RFO, Marques Jr ETA, Rodrigues LCR. 2016. Initial Description of the Presumed Congenital Zika Syndrome. Am J Public Helph 106 (4): 598–600. doi: 10.2105/AJPH.2016.303115

Nascimento EJM, Braga-Neto U, Calzavara-Silva CE, Gomes ALV, Abath FGC, Brito CAA, Cordeiro MT, Silva AM, Magalhães C, Andrade R, et al. 2009. Gene expression profiling during early acute febrile stage of dengue infection can predict the disease outcome. PloS One 4: e7892. doi: 10.1371/journal.pone.0007892

Ness RB, Haggerty CL, Harger G, Ferrell R. 2004. Differential distribution of allelic variants in cytokine genes among African Americans and White Americans. Am J Epidemiol 160: 1033–1038. doi: 10.1093/aje/kwh325

Oehler E, Watrin L, Larre P, Leparc-Goffart I, Lastere S, Valour F, Baudouin L, Mallet Hp, Musso D, Ghawche F. 2014. Zika virus infection complicated by Guillain-Barre Syndrome-case report, French Polynesia, December 2013. Euro Surveill 19: 20720. doi: 10.2807/1560-7917.ES2014.19.9.20720

Oliveira WK, França GVA, Carmo EH, Duncan BB, Kuchenbecker RS, Schimidt MI. 2017. Infection-related microcephaly after the 2015 and 2016 Zika virus outbreak in Brazil: a surveillance-based analysis. Lancet 390: 861–870. doi: 10.1016/S0140-6736(17)31368-5

Pacheco O, Beltrán, M, Nelson CA, Valencia, D, Tolosa N, Farr SL, Padilla AV, Tong VT, Cuevas EL, Espinosa-Bode A, et al. 2016. Zika Virus Disease in Colombia - Preliminary Report. N Engl J Med doi: 10.1056/NEJMoa1604037.

Parra B, Lizarazo J, Jimenez-Arango JA, Zea-Vera AF, Gonzalez-Manrique G, Vargas J, Angarita JA, Zuñiga G, Lopez-Gonzalez R, Beltran CL, et al. 2016. Guillain-Barré Syndrome associated with Zika virus infection in Colombia. N Engl J Med 375: 1513–1523. doi: 10.1056/NEJMoa1605564

Pedrosa CSG, Souza LRQ, Lima CVF, Ledur PF, Karmirian K, Gomes TA, Barbeito-Andrés J, Costa MN, Higa LM, Bellio M, et al. 2019. The cyanobacterial saxitoxin exacerbates neural cell death and brain malformations induced by Zika virus. bioRxiv doi: 10.1101/755066.

Pennington R, Gatenbee C, Kennedy B, Harpending H, Cochran G. 2009. Group differences in proneness to inflammation. Infect Genet Evol 9: 1371–1380. doi: 10.1016/j.meegid.2009.09.017

Pinto Junior VL, Lux K, Parreira R, Ferrinho P. 2015. Zika virus: a review to clinicians. Acta Med Port 28: 760–765.

Rossi AD, Faucz FR, Melo A, Pezzuto P, de Azevedo GS, Schamber-Reis BLF, Tavares JS, Mattapallil JJ, Tanuri A, Aguiar RS, Cardoso CC, et al. 2019 Variations in maternal adenylate cyclase genes are associated with congenital Zika syndrome in a cohort from Northeast, Brazil. J Intern Med 285: 215–222. doi: 10.1111/joim.12829

Salinas JL, Walteros DM, Styczynski A, Garzon F, Quijada H, Bravo E, Chaparrob P, Maderob J, Acosta-Reyesd J, Ledermann J, et al. 2017. Zika virus disease-associated Guillain-Barré syndrome-Barranquilla, Colombia 2015-2016. J Neurol Sci 381:272–277. doi: 10.1016/j.jns.2017.09.001

Santos CNO, Ribeiro DR, Alves JC, Cazzaniga RA, Magalhães LS, de Souza MSF, Fonseca ABL, Bispo AJB, Porto RLS, dos Santos CA, et al. 2019. Association between zika virus microcephaly in newborns with the rs3775291 variant in Toll-Like Receptor 3 and rs1799964 variant at Tumor Necrosis Factor-α Gene. J Inf Dis. doi: 10.1093/infdis/jiz392/5540025

Schneider WM, Chevillotte MD, Rice CM. 2014. Interferon-stimulated genes: A complex web of host defenses. Annu Rev Immunology 32: 513–545. doi: 10.1146/annurev-immunol-032713-12023

Schuler-Faccini L, Ribeiro EM, Feitosa IML, Horovitz DDG, Cavalcanti DP, Pessoa A, Doriqui MJR, Neri JI, Neto JMP, Wanderley HYC, et al. 2016. Possible association between zika virus infection and microcephaly - Brazil, 2015. MMWR 65: 59–62. doi: 10.15585/mmwr.mm6503e2

Serre D and Paabo S. 2004. Evidence for gradients of human genetic diversity within and among continents. Genome Res 14: 1679–1685.

Stephens HA. 2010. HLA and other gene associations with dengue disease severity. Curr Top Microbiol Immunol 338: 99–114.

Styczynski AR, Malta J, Krow-Lucal ER, Percio J, Nobrega ME, Vargas A, Lanzieri TM, Leite PL, Staples JE, Fischer MX, et al. 2017. Increased rates of Guillain-Barré syndrome associated with Zika virus outbreak in the Salvador metropolitan area, Brazil. PLoS Negl Trop Dis 11: e0005869. doi: 10.1371/journal.pntd.0005869

Tassaneetrithep B, Burgess TH, Granelli-Piperno A, Trumpfheller C, Finke J, Sun W, Eller MA, Pattanapanyasat K, Sarasombath S, Birx DL, et al. 2003. DC-SIGN (CD209) mediates dengue virus infection of human dendritic cells. J Exp Med 197: 823–829.

The 1000 Genomes Project Consortium. 2015. A global reference for human genetic variation. Nature 526: 68–74.

Tishkoff SA, Varkonyi R, Cahinhinan N, Abbes S, Argyropoulos G, Destro-Bisol G, Drousiotou A, Dangerfield B, Lefranc G, Loiselet J, et al. 2001. Haplotype diversity and linkage disequilibrium at human G6PD: Recent origin of alleles that confer malarial resistance. Science 293: 455–462. doi: 10.1126/science.1061573

Utgoff PE. 1989. Incremental induction of decision trees. Machine learning 4: 161–186.

Warke, RV, Xhaja K, Martin KJ, Fournier MF, Shaw SK, Brizuela N, de Bosch N, Lapointe D, Ennis FA, Rothman AL, et al. 2003. Dengue virus induces novel changes in gene expression of human umbilical vein endothelial cells. J Virol 77: 11822–11832. doi: 10.1128/jvi.77.21.11822-11832.2003

WHO. 2017. Situation report: Zika virus, microcephaly, Guillain-Barré syndrome: 20 January 2017. (Available in: http://apps.who.int/iris/bitstream/10665/254507/1/zikasitrep2Feb17-eng.pdf)

Zhong-Yu L; Wei-Feng S, Cheng-Feng Q. 2019. The evolution of zika virus from Asia to the Americas. Nature Reviews Microbiology 17:131–139.

